# Functional connectivity patterns as an early indicator of later very early preterm outcomes

**DOI:** 10.1101/2025.04.27.650837

**Authors:** Elveda Gozdas, Nehal A. Parikh, Lili He, S.M. Hadi Hosseini

## Abstract

Abnormal functional brain development associated with preterm birth has been widely reported; however, the functional brain architectures of later neurodevelopmental difficulties are not yet fully understood. Here, we applied connectome-based predictive modeling approaches to identify the brain networks associated with later neurocognitive scores at 2-3 years of age in very preterm infants (≤31 weeks’ gestation, N=79) using resting-state functional magnetic resonance imaging (rs-fMRI). The whole-brain functional connectome soon after birth successfully predicted verbal ability at 3 years of corrected age (r=0.53, 𝑝 = 4.04𝑥10^!"^) and motor ability at age 2 (r=0.39, 𝑝 = 0.0004) in very preterm infants. In particular, we found that functional edges between the frontoparietal network and limbic, motor, and medial frontal networks at birth contributed significantly to the prediction of future verbal language ability, while the edges connecting the medial frontal network and motor and basal ganglia networks contributed the most to the prediction of future motor ability. In a separate validation analysis, we demonstrated that the mean connectivity strength among these top brain networks significantly differentiated (average accuracy 76%, p<0.001) poor from normal performers at 2 and 3 years of ages. These findings highlight regional functional connectivity soon after birth as a promising biomarker for identifying risks for later brain disorders, which could inform the targeted development of effective early treatments and interventions.

## Introduction

Very preterm children are at risk for lifelong motor, cognitive, and behavioral problems (1-5). Despite advances in intensive care, more than one-third of survivors experience neurodevelopmental issues in early life and may be further diagnosed with neuropsychiatric disorders at school age, adolescence, and adulthood (6-10). Extant data suggest significant variations in the long-term outcomes of very preterm children. Preterm infants, even those with no perinatal brain abnormalities soon after birth, remain at high risk for attention deficit hyperactivity disorder, learning disabilities, language delay, and motor impairments, such as cerebral palsy (11-13). However, the field still lacks clarity on the neural characteristics in the neonatal period that can reliably predict long-term neurodevelopmental outcomes in preterm children. Advanced approaches that can identify the risk of long-term neurodevelopmental problems during the neonatal period will have significant clinical utility and can also eliminate the current practice of waiting 2–5 years to accurately evaluate neurodevelopmental difficulties in this high-risk group.

The first years of life designate the most rapid periods of brain growth and behavioral development, with complex functional pathways established in the areas responsible for language, movement, and social and emotional functioning (14-16). However, premature birth can disrupt this natural process, increasing the risk of future language, motor, and behavioral difficulties (3, 17-18). The formation of connectivity occurs quietly, and it is challenging to detect early brain disorders, such as attention deficit hyperactivity disorder (ADHD), language disorders, and cerebral palsy, which rely on behavioral changes in preterm children (10, 19-22).

Thus, defining abnormal functional connectivity using neuroimaging methods may be the best way to assess and predict subsequent neurodevelopmental disorders, as it allows the evaluation of brain structure and function during infancy.

Neuroimaging studies have provided unprecedented new insights into emerging potential abnormalities in the very preterm infant brain during infancy and offered the first compelling evidence for later neurodevelopmental disorders that reflect fundamental differences in brain structure and function (23-26). Preterm infants with suspected brain abnormalities were examined using routine structural magnetic resonance imaging (MRI). However, structural MRI is limited in its ability to detect changes in brain activity and function, such as those that occur during behavior. Thus, functional MRI can be used to provide a complete picture of brain function in preterm infants. Accumulating evidence suggests that preterm children without significant brain injury demonstrate altered functional brain activity and connectivity compared to term-born controls at term-equivalent age, several years after preterm birth, and at school age (7, 13, 27-29). Such alterations have been frequently reported in language and motor regions and regions associated with executive function and attention (13, 28). However, although this phenotype has been broadly investigated in recent years, a few studies have used neonatal neuroimaging techniques combined with advanced computational approaches (e.g., machine learning) to predict later neurodevelopmental outcomes in preterm infants (12, 28-30). The present study aimed to discover the characteristics of functional brain networks that predict long-term cognitive and motor abilities in preterm infants aged 2-3 years of age and identify functional connectivity pathways in the neonatal period that can distinguish between poor and typical longitudinal outcomes.

We conducted a longitudinal study of 79 preterm infants with neurodevelopmental outcomes at 2 and 3 years corrected ages. The hypothesis tested whether individual language and motor abilities engage specific functional brain networks and systems that may provide a neurodevelopmental marker of preterm birth, using rs-fMRI acquired during natural sleep at term-equivalent age. We employed multiple predictive modeling that can accommodate the complex interplay of whole-brain connectome and behavior to examine whether long-term language ability and cognition as measured by the DAS-II verbal, non-verbal and General Conceptual Ability (GCA), as well as and motor skills measures by the Bayley-III motor subscale can be predicted from a unique pattern of functional brain connectivity in preterm infants. We included the Bayley-III motor scores as motor function is a critical domain of early development, and impairments in motor abilities can have cascading effects on cognitive and language development. Overall, the study will uncover a unique functional brain architecture that will serve as a reliable predictor of future neurodevelopmental performance in very early preterm infants during the very early stages of their development.

## Results

### Participant Demographics and Clinical Characteristics

A cohort of 79 very preterm infants (born ≤ 31 weeks’ gestation, mean 28.52 weeks) had usable neuroimaging data (out of 101 preterm infants) at term equivalent age and long-term neurocognitive outcomes at 2-3 years of age. The demographic and clinical characteristics of the preterm infants are summarized in Table 1. The Differential Ability Scales-II (DAS-II) verbal score was used to measure the child’s ability to understand and use the language, DAS-II non-verbal language scores and GCA evaluated the cognitive skills at three years of corrected age, and the Bayley Scales of Infant and Toddler Development, third edition (Bayley-III; Motor subtest) was used to evaluate motor ability at two years of corrected age.

**Table 1.** Characteristics of the very preterm participants (n=79)

|  |  |
| --- | --- |
| Gestational age at birth in weeks (GA), mean (SD) | 28.52 (2.41) |
| Birthweight in grams, mean (SD) | 1121.15 (383.54) |
| Female sex, n (%) | 34 (42.5%) |
| Severe Bronchopulmonary dysplasia (BPD), n (%) | 20 (25%) |
| Severe Retinopathy of prematurity (ROP), n (%) | 8 (10%) |
| Postmenstrual age at MRI in weeks (PMA), mean (SD) | 40.31 (0.67) |
| DAS-II Verbal score, mean (SD) | 89.52 (21.86) |
| DAS-II Non-Verbal score, mean (SD) | 98.78 (18.8) |
| General Conceptual Ability (GCA), mean (SD) | 93.6 (18.9) |
| Bayley-III Motor composite score, mean (SD) | 94.7 (15.55) |

### Baseline functional connectivity patterns predict DAS-II neurodevelopmental verbal scores at 3 years old and Bayley-III motor scores at 2 years old in preterm infants

To define the functional network-based biomarkers of later neurodevelopmental outcomes, we analyzed rs-fMRI (motion-free, Materials, and Methods) data from seventy-nine very preterm infants (28.52±2.41, ≤31 weeks gestational age) collected at Nationwide Children’s Hospital. Whole-brain functional networks were constructed for each infant by extracting and correlating the resting-state blood oxygen level-dependent (BOLD) signal between all unique pairs of 223 functional brain regions (31). We then iteratively fitted the partial correlations between Fisher’s transformed functional connectivity values and future DAS-II neurodevelopmental Verbal and Bayley-III Motor scores controlling for demographics and clinical findings to define the connectivity features. Clusters of functional connectivity that were significantly correlated with DAS-II verbal (p <0.01) and Bayley-III (p <0.005) motor scores were retained and further separated into positive and negative correlations, referred to as positive and negative networks, respectively. These analyses revealed that 84 (90) edges were positively (negatively) correlated with DAS-II verbal scores and 68 (86) edges were positively (negatively) associated with Bayley-III motor scores. Further, principal component regression (PCR) (32) models were built using each network’s (positive and negative) edges as features with reduced components depending on the variance in each behavioral outcome using leave-one-subject-out cross-validation (LOOCV). Model prediction was assessed by the linear correlation between the predicted and observed DAS-II verbal and Bayley-III motor scores. Significant correlations emerged between the model-predicted and observed DAS-II verbal scores (r=0.53, 𝑝 = 4.04𝑥10^!"^) and Bayley-III motor scores (r=0.39, 𝑝 = 0.0004) in only positively correlated networks (Figure 2a), using bootstrap tests (1000 iterations), while the negatively correlated functional brain networks didn’t reveal the significant prediction of both language and motor development. It should be noted that these results are also significant with connectome-based predictive modeling (CPM) (33), in which averages across all edges were positively correlated with behavioral performance with lower performance (Figure S1). These results suggest that whole-brain functional connectivity predicts later neurodevelopmental outcomes in preterm infants regardless of the prediction method. It is worth mentioning that we also conducted an analysis for DAS-II Nonverbal and GCA scores, but the correlation between the observed and predicted DAS-II Non-verbal GCA scores did not reach significance for either of the models.

**Figure 1.**
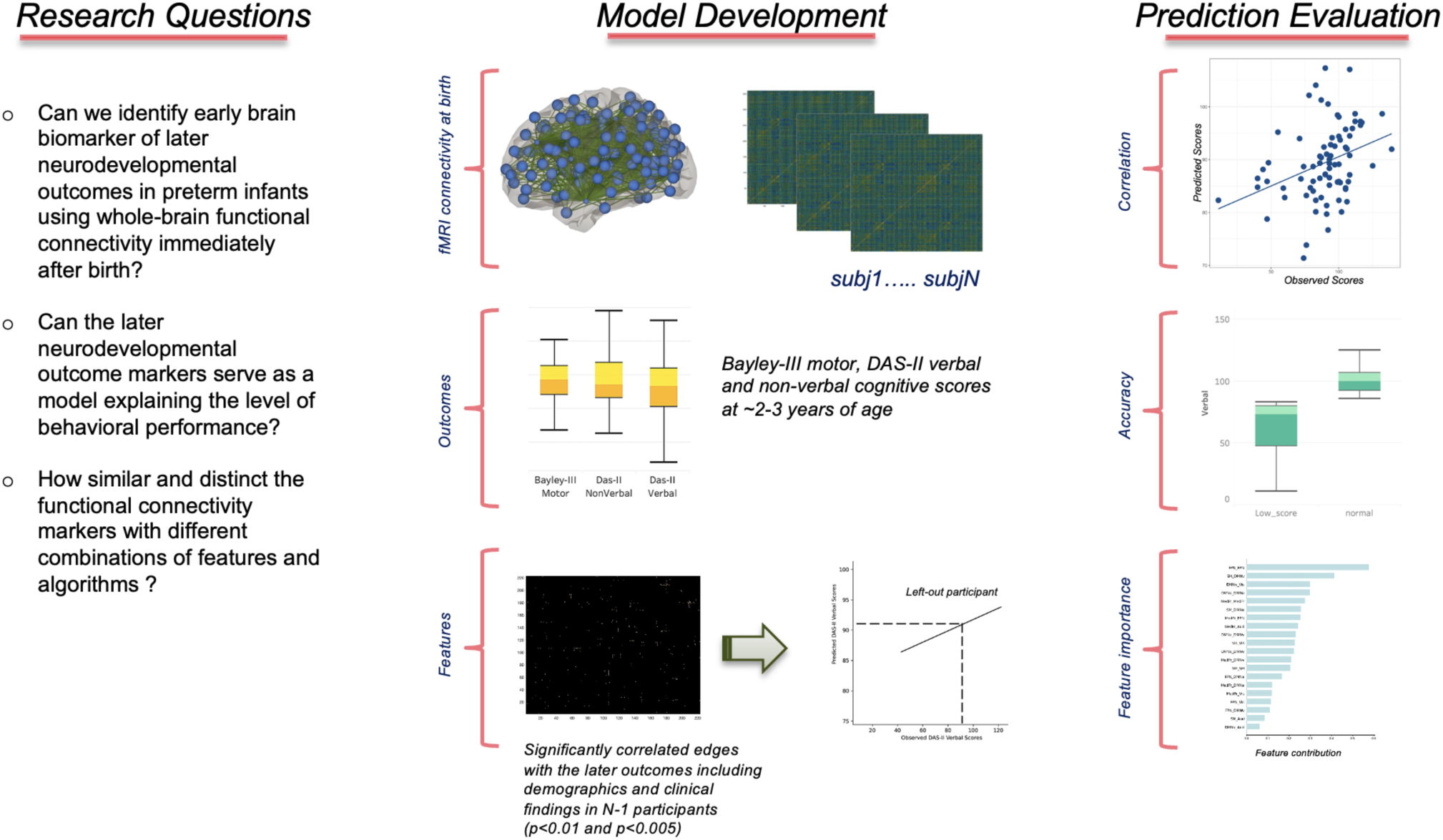
Workflow of predicting neurodevelopmental outcomes at two and three years based on resting-state functional connectivity at birth. Functional connectivity quantified from resting-state fMRI was used to predict language ability assessed with DAS-II cognitive scales at three years of age and motor ability assessed with Bayley-III Scales at two years of age. Functional connectivity edges that were significantly correlated with DAS-II verbal scores and non-verbal (p <0.01) and Bayley-III Motor scores (p <0.005) were selected as feature vectors using leave-one-out cross-validation, and prediction models were established and tested in left-out participant using principal component regression (PCR) and connectome-based predictive modeling (CMP). The correlation between predicted and actual scores evaluated prediction models. Separately, the accuracy was generated using functional connectivity among the top brain networks in categorizing the preterm infants with low and normal performers, and the feature contribution weights were quantified in the model.

**Figure 2.**
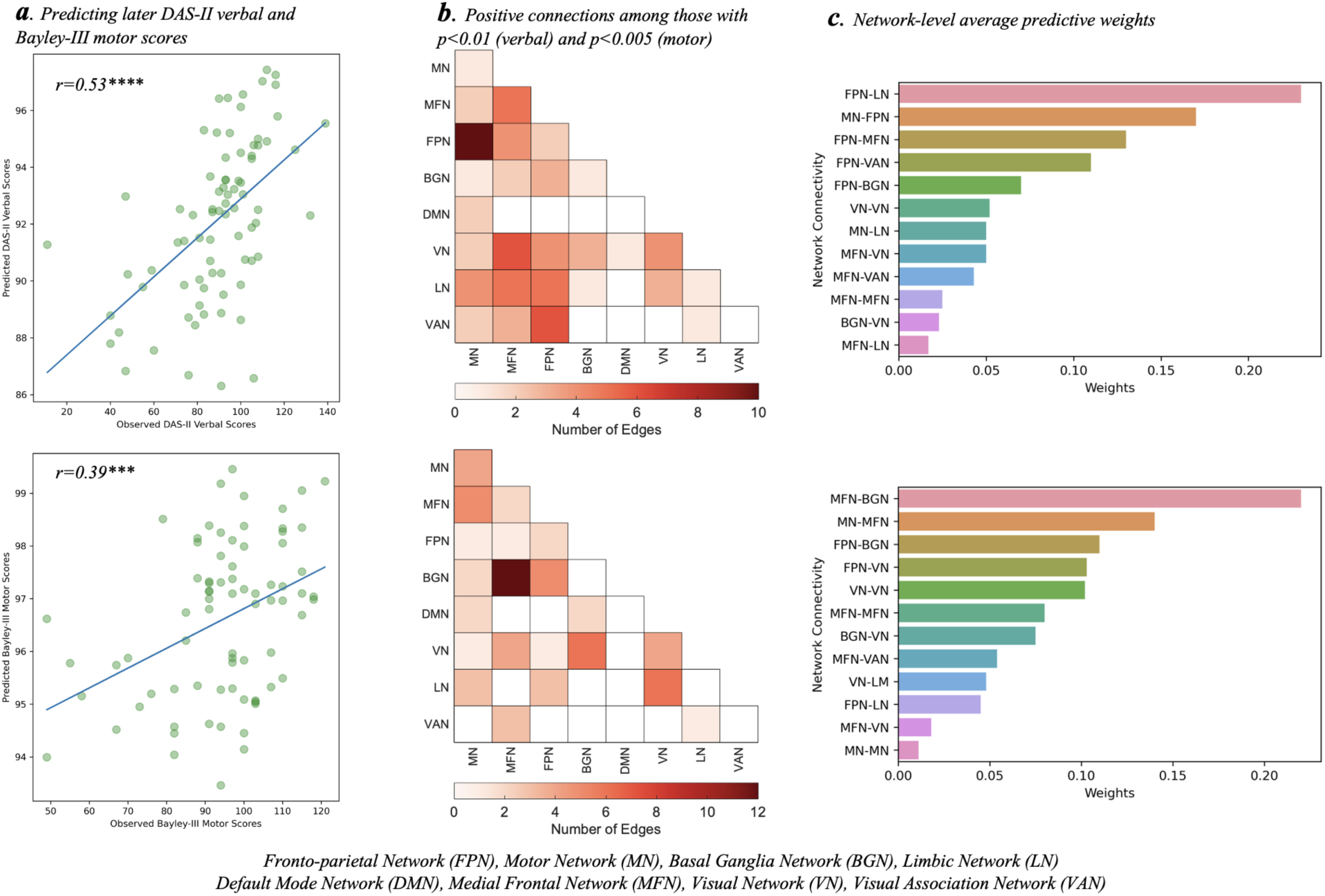
Correlation between predicted and observed DAS-II verbal scores and Bayley-III motor scores (a), The number of positive edges within and between network pairs for DAS-II verbal scores and Bayley-III motor scores (b) and their weights (c). The predicted scores were generated from positive edges with a statistical threshold (p<0.01 and p<0.005). The exact p values were 𝑝 = 4.04𝑥10^!"^ for DAS-II verbal scores and 𝑝 = 0.0004 for Bayley-III motor scores. *\*\*\*P<0.001, ****P<0.0001*.

### Network level characterization of later neurodevelopmental outcomes

To interpret the neurobiological validity of the DAS-II verbal and Bayley-III motor predictions, it is important to explore which functional brain connectivity changes were major contributors to the prediction; thus, we divided the whole brain regions into eight functional brain networks (Table S1) (33). We quantified the number of edges within and between network pairs and their weights to predict the verbal and motor outcomes (Figure 2b, c). We further conducted a bootstrap test (1000 iterations) to identify the most crucial connectivity features for prediction.

As shown in Figure 2b, the motor network (MN) and the frontoparietal network (FPN) had the highest number of positive edges that were positively correlated with DAS-II verbal scores and the strongest positive connections between the FPN and limbic network (LN), medial frontal network (MFN), and MN (Figure 2c), which seems to suggest the importance of multiple sensory and motor systems, including auditory system integration, in the language development of preterm infants. The top positive edges were mainly between the left middle temporal gyrus (MTG), left angular gyrus (ANG), and inferior parietal lobule (IPL); between the right inferior temporal gyrus (ITG) and right and left precuneus (PCUN) and right paracentral lobule (PCL); between the right ANG and left and right PCUN; and between the right orbitofrontal cortex (inf, ORBinf) and the right temporal pole (superior, TPOsup) and the left ITG, as shown in Figure 3a (blue nodes). The next most effective network pairs contributing to positive edges were between PFN-basal ganglia network (BGN) and the Visual Network (VN). In addition, multiple regression analysis showed that the mean network strengths of these networks were significantly correlated with the DAS-II verbal scores at the network level (p<0.001, r>0.7), even after correcting for multiple comparisons.

**Figure 3.**
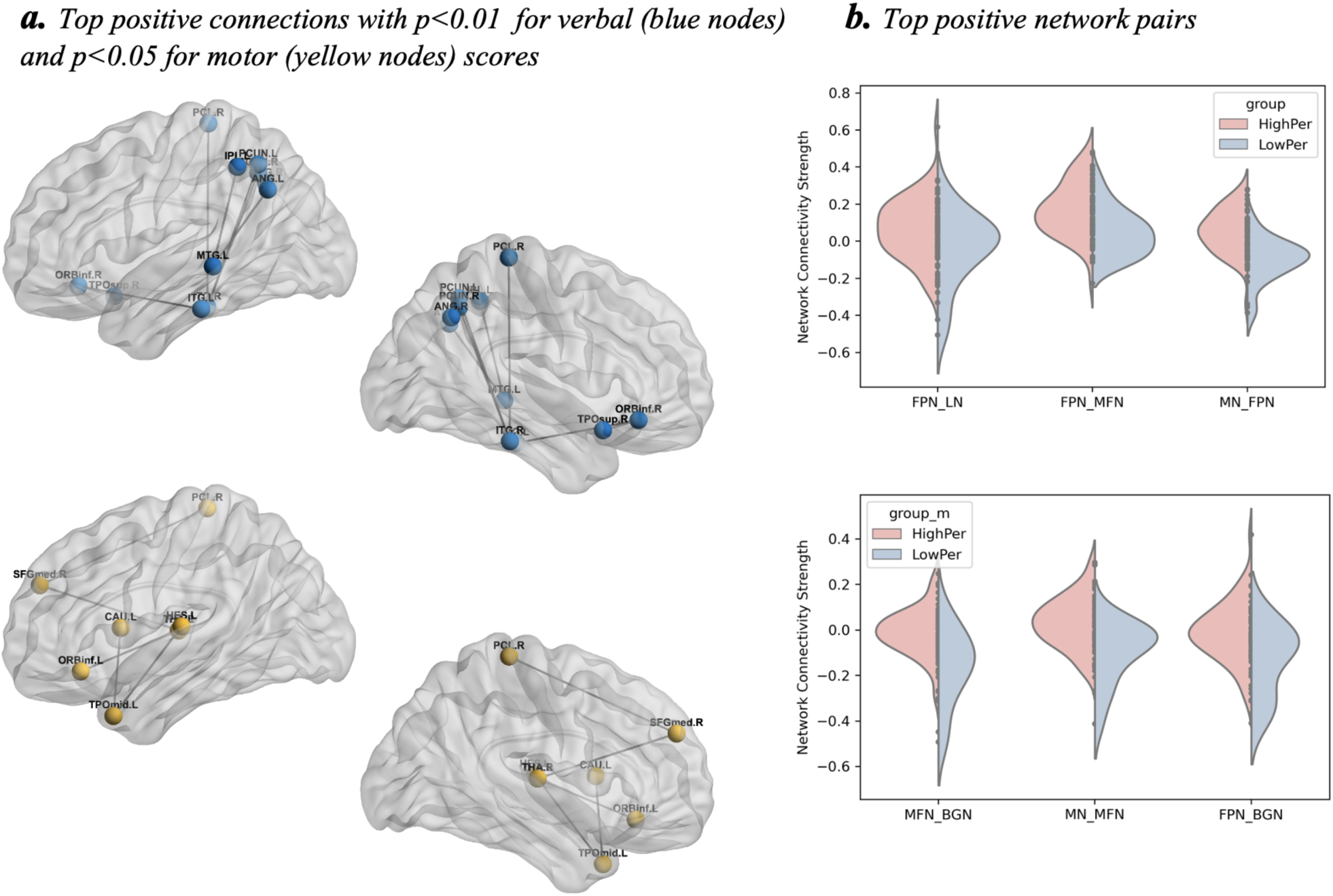
Top positive connections for predicting later DAS-II verbal and Bayley-III motor scores (a). The violin plots show the average connectivity strengths distribution between top positive network pairs that are used for classifying the low and normal verbal (top) and motor (bottom) score groups.

Similarly, the edges between the MFN, BGN, and MN contributed the most to predicting Bayley-III motor scores. Notably, the top positive edges were observed between the left thalamus (THA) and left orbitofrontal cortex (inferior, ORBinf) and temporal pole (middle, TPOmid), right superior frontal gyrus (medial, SFGmed), right THA and PCL, and left TPOmid and left Heschl gyrus (HES) and caudate (CAU), as shown in Figure 3a (yellow nodes). The mean strengths of the positive edges in these networks were also correlated with motor scores (p<0.001, r>0.63). Overall, the results demonstrated that preterm infants with higher network connectivity between the FPN and MFN, LN, and MN had higher DAS-II scores at age 3, while those with higher internetwork connectivity between the MFN and BGN and MN showed better performance in Bayley-III motor ability at age 2. Overall, the results revealed that the functional connectivity patterns for later DAS-II verbal, and Bayley-III motor scores were specific to each neurodevelopmental outcome in preterm infants.

Next, we examined whether the observed network predictors of future neurodevelopmental outcomes were explained by positive or negative functional connectivity between brain regions. The FPN network connections across the other networks, which were the most significant contributors to the DAS-II verbal score prediction, comprised mainly of positive connections (77.5% of all connections). In comparison, the MFN-BGN and MN-MFN networks, which showed the most significant contribution to predicting Bayley-III motor ability, were composed of a balanced combination of positive and negative connections (45.6% negative) during the rest.

### Evaluation of robustness of predicting the later DAS-II verbal, and Bayley-III motor scores

To further validate our findings, we tested whether functional connectivity between the identified top brain networks can accurately distinguish infants with low and normal cognitive outcomes.

Since DAS-II verbal language and Bayley-III motor scores are strong indicators of language and motor delays with specific cut-off values (34-35), the preterm infants were categorized into low (Bayley-III motor and DAS-II score ≤85) and normal (85>) groups. In addition to predicting later neurodevelopmental outcomes, the linear support vector machine (SVM) models accurately discriminated preterm children with developmental delay from children with normal development using the top three within and between positive functional network connectivity.

Similar to the previous analysis, LOOCV was used to obtain an unbiased estimate of classification performance. The average classification accuracy was 76% (AUC:74, p<0.001 binomial test) for DAS-II verbal scores and 78% (AUC: 71, p<0.001 binomial test) for Bayley-II motor scores, suggesting that network connectivity strength at term is a sensitive biomarker for future developmental delays in very preterm infants. Notably, the average functional connectivity strengths between these top network connections were significantly different between preterm children with scores>85 compared to children with developmental delay for both DAS-II verbal, and Bayley-II motor scores (𝑝_#$%_ < 0.05) (Figure 3b).

### Bayley-III motor scores predict future DAS-II verbal performance

Previous studies have shown that atypical motor development is often one of the first indications of future developmental impairments (36-37). Thus, we also evaluated whether Bayley-III motor ability at age two could predict later DAS-II verbal language skills at age 3 in preterm children. Using a multiple linear regression model, we observed that Bayley-III motor scores significantly predicted later DAS-II verbal ability (p=0.005, r=0.4), corroborating the idea that the presence of motor delays is often associated with developmental delays in other domains in very preterm children.

## Discussion

In this study, using recently developed computational approaches, we built functional network models that carry predictive information for later neurodevelopmental outcomes measured at 2-3 years of age in preterm infants based on whole-brain functional connectivity. We discovered patterns of functional brain connectivity that were predictive of later neurodevelopmental outcomes. Specifically, functional connectivity between the FPN network and other brain networks predicted later language development, and connectivity between the MFN, BGN, and MN predicted later motor ability in preterm infants. In addition, the identified top functional brain networks that predicted later language and motor outcomes significantly discriminated between preterm children with typical development and those with developmental delays. Our findings provide valuable insights into how later neurodevelopmental difficulties can be predicted from a preterm infant’s unique functional brain connectivity at birth.

With the apparent progress in the survival rate of very preterm infants, there is an increasing focus on characterizing later preterm neurodevelopmental problems during the neonatal period. It has been well documented that very preterm birth is associated with learning, academic achievement, behavioral abnormalities, and language issues (38-41). Although previous neuroimaging studies have extensively reported disruptions in functional connectivity across brain regions important for language function in very preterm infants (13, 42–44), few studies have focused on predicting developmental outcomes. Here, we explored whether the functional connectivity patterns are crucial to language development in very preterm infants. Notably, we found that the connections between FPN, MFN, MN, and LN were among the most important predictors of language development in preterm infants. Language function is intricately linked to other higher cognitive processes beyond its fundamental role in communication (45), and these findings highlight the importance of the interconnections between the FPN, and other networks related to other functions such as memory and cognitive processes predicting later language problems in very preterm infants during infancy. Further, the top positive edges were represented on the right and left, which also explains how reduced lateralization on the left and increased connectivity on the right (46) are associated with future language development.

Abnormalities in various functional brain areas, similar to preterm infants with periventricular leukomalacia and severe intraventricular hemorrhage, may contribute to motor problems, such as cerebral palsy, and children may develop motor problems even with no high-grade brain injury.(47). Developmental motor delays and widespread disruptions of functional connectivity in motor-related brain areas have been well characterized in very preterm infants, school-aged children, and adults born prematurely (13, 43, 48). Similar to other neurodevelopmental outcomes, there is limited research investigating the association between functional connectivity at birth and later motor development in preterm infants (30, 47, 49). The study showed that the functional edges between the MFN-BGN and MN-MFN networks during the neonatal period effectively predicted motor performance at two years. The internetwork connections that strongly contribute to the prediction of later motor performance are important for motor function and are largely consistent with previous findings reporting abnormal changes in these connections in preterm infants (44, 50). Furthermore, primary and motor brain regions exhibit earlier maturation and connect with other brain regions to build key parts of an infant’s brain development (51).

Thus, motor outcomes could serve as early biomarkers for later neurodevelopment in preterm infants. In line with recent research, we also found that two-year motor skills predicted language ability at three years old, suggesting that early disruptions in motor functioning are predictive of future impairments in language development in very preterm infants.

The current study provides a neuroimaging-based biomarker for later neurodevelopmental problems in very preterm infants to diagnose and evaluate brain disorders at birth. The extent to which brain representation at birth overlaps with subsequent neurodevelopmental problems remains an unsolved question. Here, we found that whole-brain functional network models classified children with aberrant neurodevelopment from typical development with a reasonable average accuracy level. Examination of the models’ anatomical representation showed that the functional brain network patterns were largely crucial for language and motor function, but the models also identified some connections that have not been well documented, such as the positive weights between the left TPOmid and left Heschl gyrus (HES) for motor prediction (52).

Therefore, this study provides an interesting future direction for examining the different functional roles of these connections in language and motor development in preterm infants.

This study had some limitations that should be acknowledged. The preterm infants were scanned over a two-year period, and the fMRI scanning protocol was upgraded to multiband image acquisition to shorten the acquisition time. This resulted in a range of rs-fMRI time-points, which may have led to inconsistencies. To address this, we matched the same number of time points for both multiband and non-multiband fMRI data, the distribution of subjects with multiband and non-multiband fMRI data was very similar in the prediction model in each validation fold.

Furthermore, we were unable to test the validity of the models in an independent sample due to difficulties with neonatal neuroimaging data at birth with later neurodevelopmental outcomes in preterm infants. However, we tested the robustness of our models with linear SVM analysis, which showed a very high performance for classifying the low and normal development groups in preterm infants. Finally, our study didn’t reveal significant findings for DAS-II nonverbal and GCA score predictions that may reflect these higher-order cognitive abilities are supported by distributed brain networks that are not yet fully mature at term-equivalent age. Furthermore, later cognitive outcomes are likely influenced by a combination of brain development and postnatal environmental factors that are not fully captured by early functional connectivity patterns. Future studies with longitudinal imaging and environmental assessments will be important to better understand how early brain network development interacts with postnatal experiences to shape later nonverbal language and general cognitive abilities.

In conclusion, we identified that functional brain networks at birth provide insights into the neurobiological mechanisms that support later language and motor ability components in preterm infants. Crucially, the current study facilitated the discovery of these functional network weights for predicting later language and motor performance by considering whole-brain connections rather than limiting them to specifying a priori connections. Therefore, these findings advance the understanding of future neurodevelopmental problems in preterm infants, which could have implications for the early diagnosis of brain disorders and interventions around birth.

## Materials and Methods

### Participants

The total sample consisted of 110 very preterm infants (≤31 weeks gestational age) recruited from the Nationwide Children’s Hospital, Ohio State University Medical Center, and two affiliated level III NICUs from Columbus, Ohio (34 girls, mean gestation=28.52, SD=2.41). Infants with known structural congenital central nervous system anomalies, congenital chromosomal anomalies, congenital cyanotic cardiac defects, visible defects in the white or grey matter, and excessive motion were excluded. We included 79 very preterm infants without significant brain injury on anatomical MRI. The Nationwide Children’s Hospital Institutional Review Board approved this study and written informed consent was obtained from all subjects prior to imaging.

### Later neurodevelopmental outcomes and clinical assessments

The assessment of neurodevelopmental outcomes for very early preterm infants was completed using the Differential Ability Scale, Second Edition (DAS-II, Verbal score) (53) at three years of corrected age and Bayley Scales of Infant and Toddler Development, third edition (Bayley-III; Motor composite score) (54) at two years of corrected age. The DAS-II is an individually administered test designed to measure distinct cognitive abilities in children and adolescents ages two years, six months to 17 years. The DAS-II comprises individual subtests (verbal and non-verbal reasoning and spatial skills) that evaluate the strengths and weaknesses of a broad range of learning processes. The Verbal Scale assesses children’s language comprehension and usage abilities, including vocabulary, grammar, and verbal reasoning. It is one of the scales included in the DAS-II assessment and used by psychologists, educators, and other specialists to diagnose learning disabilities, monitor progress, and plan interventions. The Bayley-III motor subscale is a commonly used standardized assessment tool to evaluate a child’s gross and fine motor skills, including controlling movement and balance and performing age-appropriate motor activities.

Baseline demographics, clinical outcome data, later DAS-II Verbal and non-verbal scores, and Bayley-III Motor scores are included in Table 1.

### MRI Data and Acquisition and Preprocessing

#### MRI Data Acquisition

Preterm infants (mean postmenstrual age at scan 39.8 weeks, range 39-41 weeks) were scanned on a 3T Siemens Skyra scanner using a 32-channel head coil during natural sleep. High-resolution anatomical T2-weighted images were acquired to align the fMRI data and inspect the brain anatomy. T2-weighted images were obtained with TE=147ms, TR=9500ms, 0.93×0.93×1.1 mm voxel size, flip angle=150°, and FOV=180×180 mm. The rs-fMRI data were collected from a gradient echo-planar sequence with TE=30ms, TR=2700ms, FOV=212×212 mm, matrix size=104×104, slice thickness 2.5 mm) and 200 measurements in 38 preterm infants. In addition, multiband rs-fMRI data were acquired from 41 preterm infants with TE=30ms, TR=1000ms, FOV=212×212 mm, matrix size=104×104, slice thickness 2.5 mm, and 500 measurements.

#### fMRI Data Preprocessing

SPM (https://www.fil.ion.ucl.ac.uk/spm/) was used to preprocess the rs-FMRI data. The fMRI data were corrected for head motion by realigning all images to the mean of all functional volumes. Although the neonates were asleep during the fMRI procedures, head motion was still problematic and was corrected, as described in detail in the section below. The mean functional image was co-registered with the corresponding T2-weighted high-resolution image. T2-weighted images were segmented into gray matter, white matter, and cerebrospinal fluid (CSF) using neonatal tissue probability maps (55). Tools developed for segmenting adult brain images also do not work in neonatal brain images because of the reduced contrast between gray matter and white matter that arises from the higher water content and limited myelination in the neonatal brain. Consequently, we utilized a neonatal brain atlas explicitly developed to address this low-contrast condition by introducing age-appropriate prior probability estimates (55). Finally, the images were normalized to the neonatal template (56), re-sliced to a voxel size of 2×2×2 mm, and smoothed with a full width at half maximum (FWHM) Gaussian Smoothing kernel of 6 mm. T1-weighted images in neonates did not provide sufficient contrast for segmentation and co-registration, and although acquired, they were not used in our analysis.

#### Functional Network Construction

Whole-brain functional connectivity for pre-processed rs-FMRI data was performed using the Conn functional connectivity toolbox (57). Conn is a MATLAB-based toolbox for computational analysis of functional connectivity in fMRI (fcMRI). The residual BOLD time series was extracted from the gray matter voxels. The rs-fMRI time series from each voxel was corrected using a strict noise reduction method called aCompCor, which removed the principal components attributed to white matter and cerebrospinal fluid signals (58) and eliminated the need for global signal regression (59-60). In addition, six subject-specific motion parameter time series and their first derivatives and extreme outlier volumes detected with a threshold of 0.5 mm in ART (http://www.nitrc.org/ projects/artifact_detect/) were introduced as potential confounders. ART is a graphic tool for automatic and manual detection of global mean and motion outliers in fMRI data. In addition, the residual BOLD time series in each voxel was band-pass filtered at 0.008 to 0.08 Hz to focus on low-frequency fluctuations. Next, whole-brain networks were computed for each participant using a recently generated neonatal functional brain atlas consisting of 223 regions of interest (ROIs) (31) (Table S2). Given our focus on functional connectivity analyses, we used the neonatal functional brain atlas as it provides a more precise mapping of functional networks than structural atlases based on anatomical landmarks (31). The BOLD signal was extracted from each ROI during rest and bivariate correlations were computed between each pair of ROIs, resulting in a 223 × 223 correlation matrix for each participant.

### Whole-brain Connectome-modelling

We adopted principal component regression (PCR) with a reduced number of PCs and connectome-based predictive modeling (CPM) methods to determine whether resting-state functional connectivity can be applied to predict later neurodevelopmental performance in very preterm infants. First, we identified the features for each neurodevelopmental outcome correlating each edge’s strength in resting-state functional connectivity with neurodevelopmental scores using different types of correlation methods (linear and partial correlations including clinical and demographic data including gestational age, birth weight, sex, as well as BPD, and ROP due to their strong associations with neonatal brain development and outcomes. Correlations were additionally controlled for mean displacement to minimize the potential confounding effects of head motion. Next, we defined positive and negative networks that showed positive and negative correlations with the verbal, non-verbal, and GCA or motor scores with multiple levels of thresholding on p values of correlations (p<0.05-0.001), but the threshold at which significant prediction was achieved varied across outcomes: language development required p < 0.01, while motor development was predicted at p < 0.005. Leave-one-out cross-validation (LOOCV) was used to obtain unbiased results. We used LOOCV due to the limited sample size, which allowed for maximal use of the available data in training the predictive models. The predictive model was then applied to one infant’s data by summing the edge strengths in the positive and negative networks to generate the predicted scores. The positive edges were input into the PCR as features, whereas the average of all positive edges was input into the CPM as a feature. A linear regression model was used to estimate the relationship between model-predicted and observed neurodevelopmental scores. It should be noted that the positive edges selected using the partial correlation with the thresholds p<0.01 for verbal and p<0.005 for motor scores yielded significant correlations between observed and predicted scores, whereas the other correlation thresholds selecting the functional connectivity edges, including negative edges, did not provide significant results between predicted and observed.

### Evaluation of model performance for predicting later neurodevelopmental outcomes

We first calculated the linear correlation values between the actual and observed later neurodevelopmental outcomes and averaged the correlation values to obtain one value per outcome score using 1000 iterations. Next, to validate the performance of connectivity strength in verbal and motor score predictions, the DAS-II verbal, and Bayley-III motor scores were categorized as normal (85>) and low (85≤) scores. Using the linear SVM model, including demographics and clinical findings, positive edges were used as features to classify each infant’s score into normal and low scores. We also used LOOCV, and the average accuracy was used as the classification accuracy. The area under the receiver operating characteristic (ROC) curve was used to evaluate the performance of the classification model. Finally, the binomial tests were applied to determine whether the classification accuracies were significantly better than the distribution of expected classification results, with a 50% chance-level probability.

Finally, to identify the functional pattern of the verbal and motor networks, we classified the infant functional brain atlas into eight functional networks (FPN, MN, BGN, LN, DMN, MFN, VN, and VAN; Table S1) (31, 33). We next discovered the characteristics of within- and between-network connectivity by extracting the number of edges between and within defined networks and conducted bootstrap tests (with 1000 iterations) to identify the connectivity features that contributed the most to the prediction using a gradient boosting algorithm and grouping the weights into the brain network regions.

## Acknowledgements

We sincerely thank Wells Logan, MD, and Kelly McNally, PhD, for supervising the neurodevelopmental testing; Jennifer Notestine, RN and Valerie Marburger, NNP, for serving as the study coordinators; and Mark Smith, MS, for serving as the study MR technologist. We are also grateful to the families and Biobehavioral Core staff that made this study possible. This work was supported by the National Institute of Neurological Disorders and Stroke of the National Institutes of Health [grant numbers R01-NS094200 and R01-NS096037]. EG’s efforts were supported by the Eunice Kennedy Shriver National Institute of Child Health and Human Development [grant number K99HD109507] and the Stanford Maternal and Child Health Research Institute (MCHRI). The content is the sole responsibility of the authors and does not necessarily represent the official views of the National Institutes of Health.

## Data Access Statement

Research data supporting this publication are not publicly available due to the inclusion of sensitive information within the dataset.

## Author Contributions

EG carried out the image processing and statistical analysis and drafted the manuscript. NNP supervised the project and contributed to MRI data collection, the research design and implementation, and the manuscript’s drafting. LH contributed to drafting the manuscript. SMH also supervised the project and helped drafting the manuscript. All authors read and approved the final manuscript.

## Supplemental Information

**Figure S1.**
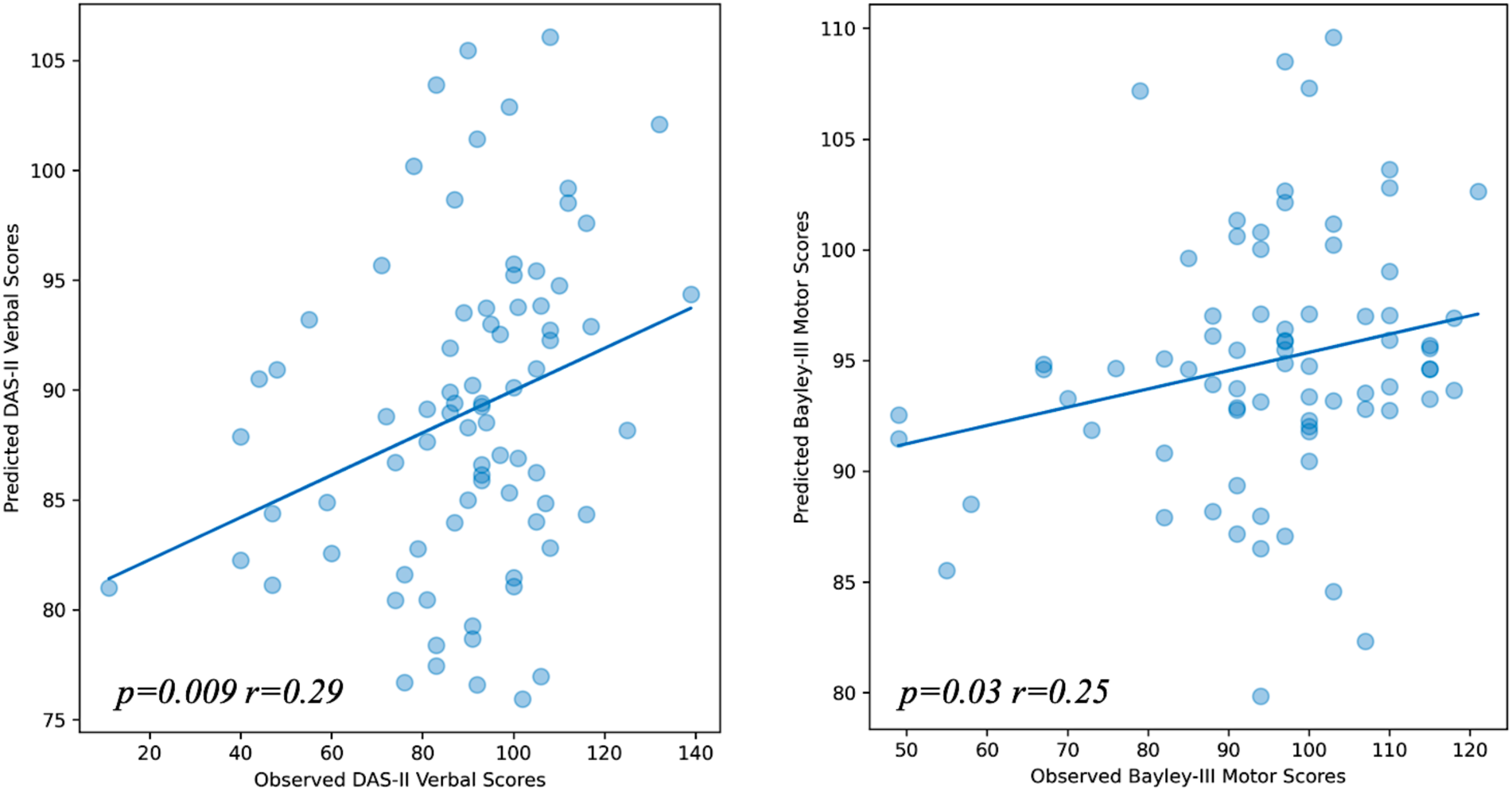
Correlation between predicted and observed DAS-II Verbal scores (a) Correlation between predicted and observed Bayley-III Motor scores (b) using connectome-based predictive modelling. The predicted scores are generated from positive edges with a statistical threshold (p<0.01).

**Table S1.** Table of regions of interest (ROIs) in the neonatal functional brain atlas (Shi et al., 2018) and corresponding brain networks.

| *Fronto-parietal Network*<br>*(FPN)* | *Medial Frontal Network*<br>*(MFN)* | *Motor Network (MN)* | *Default Mode Network (DMN)* |
| --- | --- | --- | --- |
| Orbitofrontal cortex (superior) | Superior frontal gyrus (dorsal) | Precentral gyrus | Orbitofrontal cortex (medial) |
| Middle frontal gyrus | Inferior frontal gyrus (opercular) | Rolandic operculum | Posterior cingulate gyrus |
| Orbitofrontal cortex (middle) | Orbitofrontal cortex (inferior) | Supplementary motor area |  |
| Inferior frontal gyrus (triangular) | Superior frontal gyrus (medial) | Paracentral lobule |  |
| Inferior parietal lobule | Rectus gyrus | Postcentral gyrus |  |
| Angular gyrus | Anterior cingulate gyrus | Supramarginal gyrus |  |
| Inferior temporal gyrus | Middle temporal gyrus |  |  |
|  | Temporal pole (middle) | Heschl gyrus |  |
|  |  | Superior temporal gyrus |  |
|  |  | Temporal pole (superior) |  |
| *Basal Ganglia Network*<br>*(BGN)* | *Limbic Network (LN)* | *Visual Network (VN)* | *Visual Association Network*<br>*(VAN)* |
| Olfactory | Insula | Calcarine cortex | Middle occipital gyrus |
| Hippocampus | Middle cingulate gyrus | Cuneus | Inferior occipital gyrus |
| ParaHippocampal gyrus | Precuneus | Lingual gyrus | Superior parietal gyrus |
| Caudate | Amygdala | Superior occipital gyrus |  |
| Putamen |  | Fusiform gyrus |  |
| Thalamus |  |  |  |
| Pallidum |  |  |  |

**Table S2.** The ROIs defined in the neonatal functional brain atlas.

| Regions | Name | Regions | Name | Regions | Name | Regions | Name |
| --- | --- | --- | --- | --- | --- | --- | --- |
| 1 | Precentral gyrus left | 61 | Olfactory right | 121 | Lingual gyrus right | 181 | Thalamus right |
| 2 | Precentral gyrus left | 62 | Superior frontal gyrus (medial) left | 122 | Superior occipital gyrus left | 182 | Thalamus right |
| 3 | Precentral gyrus right | 63 | Superior frontal gyrus (medial) right | 123 | Superior occipital gyrus left | 183 | Heschl gyrus left |
| 4 | Precentral gyrus right | 64 | Superior frontal gyrus (medial) right | 124 | Superior occipital gyrus right | 184 | Heschl gyrus left |
| 5 | Precentral gyrus right | 65 | Superior frontal gyrus (medial) right | 125 | Superior occipital gyrus right | 185 | Heschl gyrus right |
| 6 | Superior frontal gyrus (dorsal) left | 66 | Superior frontal gyrus (medial) right | 126 | Middle occipital gyrus left | 186 | Superior temporal gyrus left |
| 7 | Superior frontal gyrus (dorsal) left | 67 | Superior frontal gyrus (medial) right | 127 | Middle occipital gyrus left | 187 | Superior temporal gyrus right |
| 8 | Superior frontal gyrus (dorsal) left | 68 | Orbitofrontal cortex (medial) left | 128 | Middle occipital gyrus right | 188 | Superior temporal gyrus right |
| 9 | Superior frontal gyrus (dorsal) left | 69 | Orbitofrontal cortex (medial) right | 129 | Middle occipital gyrus right | 189 | Superior temporal gyrus right |
| 10 | Superior frontal gyrus (dorsal) left | 70 | Orbitofrontal cortex (medial) right | 130 | Inferior occipital gyrus left | 190 | Superior temporal gyrus right |
| 11 | Superior frontal gyrus (dorsal) left | 71 | Orbitofrontal cortex (medial) right | 131 | Inferior occipital gyrus right | 191 | Superior temporal gyrus right |
| 12 | Superior frontal gyrus (dorsal) left | 72 | Rectus gyrus left | 132 | Fusiform gyrus left | 192 | Superior temporal gyrus right |
| 13 | Superior frontal gyrus (dorsal) left | 73 | Rectus gyrus left | 133 | Fusiform gyrus left | 193 | Superior temporal gyrus right |
| 14 | Superior frontal gyrus (dorsal) left | 74 | Rectus gyrus left | 134 | Fusiform gyrus left | 194 | Temporal pole (superior) left |
| 15 | Superior frontal gyrus (dorsal) left | 75 | Rectus gyrus left | 135 | Fusiform gyrus left | 195 | Temporal pole (superior) left |
| 16 | Superior frontal gyrus (dorsal) right | 76 | Rectus gyrus left | 136 | Fusiform gyrus left | 196 | Temporal pole (superior) left |
| 17 | Superior frontal gyrus (dorsal) right | 77 | Rectus gyrus left | 137 | Fusiform gyrus left | 197 | Temporal pole (superior) left |
| 18 | Superior frontal gyrus (dorsal) right | 78 | Rectus gyrus right | 138 | Fusiform gyrus left | 198 | Temporal pole (superior) right |
| 19 | Superior frontal gyrus (dorsal) right | 79 | Rectus gyrus right | 139 | Fusiform gyrus right | 199 | Temporal pole (superior) right |
| 20 | Superior frontal gyrus (dorsal) right | 80 | Rectus gyrus right | 140 | Fusiform gyrus right | 200 | Temporal pole (superior) right |
| 21 | Superior frontal gyrus (dorsal) right | 81 | Rectus gyrus right | 141 | Fusiform gyrus right | 201 | Temporal pole (superior) right |
| 22 | Superior frontal gyrus (dorsal) right | 82 | Rectus gyrus right | 142 | Fusiform gyrus right | 202 | Temporal pole (superior) right |
| 23 | Superior frontal gyrus (dorsal) right | 83 | Rectus gyrus right | 143 | Fusiform gyrus right | 203 | Middle temporal gyrus left |
| 24 | Superior frontal gyrus (dorsal) right | 84 | Insula left | 144 | Fusiform gyrus right | 204 | Middle temporal gyrus left |
| 25 | Superior frontal gyrus (dorsal) right | 85 | Insula left | 145 | Postcentral gyrus left | 205 | Middle temporal gyrus right |
| 26 | Superior frontal gyrus (dorsal) right | 86 | Insula right | 146 | Postcentral gyrus right | 206 | Middle temporal gyrus right |
| 27 | Superior frontal gyrus (dorsal) right | 87 | Insula right | 147 | Superior parietal gyrus left | 207 | Temporal pole (middle) left |
| 28 | Orbitofrontal cortex (superior) right | 88 | Anterior cingulate gyrus left | 148 | Superior parietal gyrus left | 208 | Temporal pole (middle) left |
| 29 | Orbitofrontal cortex (superior) right | 89 | Anterior cingulate gyrus left | 149 | Superior parietal gyrus right | 209 | Temporal pole (middle) left |
| 30 | Orbitofrontal cortex (superior) right | 90 | Anterior cingulate gyrus right | 150 | Superior parietal gyrus right | 210 | Temporal pole (middle) right |
| 31 | Orbitofrontal cortex (superior) right | 91 | Middle cingulate gyrus left | 151 | Inferior parietal lobule left | 211 | Inferior temporal gyrus left |
| 32 | Orbitofrontal cortex (superior) right | 92 | Middle cingulate gyrus left | 152 | Inferior parietal lobule left | 212 | Inferior temporal gyrus left |
| 33 | Middle frontal gyrus left | 93 | Middle cingulate gyrus right | 153 | Inferior parietal lobule left | 213 | Inferior temporal gyrus left |
| 34 | Middle frontal gyrus left | 94 | Middle cingulate gyrus right | 154 | Inferior parietal lobule left | 214 | Inferior temporal gyrus left |
| 35 | Middle frontal gyrus right | 95 | Middle cingulate gyrus right | 155 | Inferior parietal lobule right | 215 | Inferior temporal gyrus left |
| 36 | Middle frontal gyrus right | 96 | Middle cingulate gyrus right | 156 | Inferior parietal lobule right | 216 | Inferior temporal gyrus right |
| 37 | Middle frontal gyrus right | 97 | Posterior cingulate gyrus left | 157 | Supramarginal gyrus left | 217 | Inferior temporal gyrus right |
| 38 | Middle frontal gyrus right | 98 | Posterior cingulate gyrus right | 158 | Supramarginal gyrus left | 218 | Inferior temporal gyrus right |
| 39 | Orbitofrontal cortex (middle) left | 99 | Hippocampus left | 159 | Supramarginal gyrus left | 219 | Inferior temporal gyrus right |
| 40 | Orbitofrontal cortex (middle) left | 100 | Hippocampus right | 160 | Supramarginal gyrus right | 220 | Inferior temporal gyrus right |
| 41 | Orbitofrontal cortex (middle) left | 101 | Hippocampus right | 161 | Angular gyrus left | 221 | Inferior temporal gyrus right |
| 42 | Orbitofrontal cortex (middle) right | 102 | ParaHippocampal gyrus left | 162 | Angular gyrus right | 222 | Inferior temporal gyrus right |
| 43 | Orbitofrontal cortex (middle) right | 103 | ParaHippocampal gyrus left | 163 | Angular gyrus right | 223 | Inferior temporal gyrus right |
| 44 | Orbitofrontal cortex (middle) right | 104 | ParaHippocampal gyrus left | 164 | Angular gyrus right |  |  |
| 45 | Inferior frontal gyrus (opercular) left | 105 | ParaHippocampal gyrus left | 165 | Precuneus left |  |  |
| 46 | Inferior frontal gyrus (opercular) right | 106 | ParaHippocampal gyrus right | 166 | Precuneus left |  |  |
| 47 | Inferior frontal gyrus (triangular) left | 107 | ParaHippocampal gyrus right | 167 | Precuneus right |  |  |
| 48 | Inferior frontal gyrus (triangular) right | 108 | Amygdala left | 168 | Precuneus right |  |  |
| 49 | Inferior frontal gyrus (triangular) right | 109 | Amygdala right | 169 | Paracentral lobule left |  |  |
| 50 | Orbitofrontal cortex (inferior) left | 110 | Calcarine cortex left | 170 | Paracentral lobule right |  |  |
| 51 | Orbitofrontal cortex (inferior) right | 111 | Calcarine cortex right | 171 | Caudate left |  |  |
| 52 | Orbitofrontal cortex (inferior) right | 112 | Calcarine cortex right | 172 | Caudate left |  |  |
| 53 | Rolandic operculum left | 113 | Calcarine cortex right | 173 | Caudate right |  |  |
| 54 | Rolandic operculum right | 114 | Cuneus left | 174 | Caudate right |  |  |
| 55 | Supplementary motor area left | 115 | Cuneus left | 175 | Caudate right |  |  |
| 56 | Supplementary motor area left | 116 | Cuneus right | 176 | Putamen left |  |  |
| 57 | Supplementary motor area right | 117 | Cuneus right | 177 | Putamen right |  |  |
| 58 | Supplementary motor area right | 118 | Lingual gyrus left | 178 | Pallidum left |  |  |
| 59 | Olfactory left | 119 | Lingual gyrus left | 179 | Pallidum right |  |  |
| 60 | Olfactory right | 120 | Lingual gyrus right | 180 | Thalamus left |  |  |

